# Language impairment resulting from a de novo deletion of 7q32.1-q33: a case report

**DOI:** 10.1101/047241

**Authors:** Ma Salud Jiménez-Romero, Montserrat Barcos-Martínez, Isabel Espejo-Portero, Antonio Benítez-Burraco

## Abstract

Chromosome 7 is a hot spot for cognitive disorders involving language deficits. We report on a girl who presents with a cognitive and speech delay, motor problems, hearing loss, and behavioral disturbances, and a de novo deletion within 7q32.1-q33 (chromosome position: chr7:127109685-132492196, hg 18). Several genes involved in brain development and function are located within the deleted region. Many of them are related to developmental disorders encompassing language deficits (dyslexia, speech-sound disorder, and autism). The proband’s phenotype may result from a change in the expression level of some of these genes.

## INTRODUCTION

Developmental disorders entailing language deficits provide with crucial evidence of the molecular and genetic underpinnings of the human faculty for language. In the last decades, many candidate genes for disorders like dyslexia, specific language impairment (SLI), or speech-sound disorder (SSD) have been identified and functionally characterised (reviewed in ^1^). Rare or sporadic conditions resulting from chromosomal rearrangement or copy-number variation provide with additional evidence of genetic factors that are important for the development and function of the brain areas involved in language processing. Interestingly, many of them are found in chromosome 7. Specifically, the region 7q31-q36 has been claimed to represent a hot spot for the evolution of human-specific communication abilities ^2^. Within this region one finds *FOXP2* (located in 7q31), which is important for the development of brain circuits involved in procedural learning and which is found mutated in people with speech and language problems ^3,4^. The region has been related as well to disorders involving phonological deficits, like dyslexia or speech-sound disorder ^5,6^, and to autism ^7,8^. In this paper, we report on a girl who presents with a language disorder, cognitive delay, motor problems, hearing loss and behavioral disturbances, and a de novo deletion within the region 7q32.1-q33. Given the phenotype of this patient, we hypothesize that some gene(s) important for language development may have been affected by the deletion.

## CASE PRESENTATION

### Clinical History

The patient was born after 41 weeks of gestation to a 28 year old female. The proband’s parents were healthy and non-consanguineous. During the pregnancy, intrauterine growth restriction (IUGR) was reported. At birth, her weight was 2.545 kg, her height 50 cm and the cephalic perimeter 33 cm. APGAR evaluation scores were normal (8/9). Cord blood pH was 7.30. Dystocia and fetal distress were reported at delivery. Because of skin pallor and mild respiratory distress at birth, the newborn was kept in an incubator for 8 days. Further exploration was indicative of hypotonia, sucking weakness, and absence of crying (the child began crying only at 3 years). Ophthalmologic examination and cranial magnetic resonance imaging (MRI), performed at 3 years, rendered normal, although mild microcephaly was reported. The child has shown no sign of epilepsy. At 3;6 years she was diagnosed with severe bilateral sensorineural hearing loss (neuropathy). Because her medical history was significant for numerous episodes of serous otitis media, ear drainage tubes were placed. From age 4 until present she has used behind-the-ear (BTE) hearing aid. An audiometry (Auditory Steady State Response, ASSR) performed at 7 years revealed hearing loss in both ears (right ear: objective threshold of 47 dB correlated with less than 10 dB on subjective thresholds; left ear: objective threshold of 72 dB correlated with less than 10 dB on subjective thresholds). Mouth, tongue, teeth and palate examination performed at 7;7 years revealed no abnormalities.

### Language and cognitive development

Early developmental milestones were achieved normally by the child. She was able to sit without aid only at 8 months and walked at 1;6 years. Nonetheless, at age 2 her parents reported absence of speech and lack of interest in her social environment. In order to accurately evaluate her global development, the Spanish version of the Battelle Developmental Inventories ^9^ was administered at ages 2, 3 5, and 7. The obtained scores pointed to a broad developmental delay, which has exacerbated over the years and that impacts language mostly. Accordingly, communication skills were severely impaired; cognitive and personal-social abilities were impaired, and adaptive abilities and motor skills were relatively spared (figure 1). From age 3 onwards the proband has exhibited impulsive behavior. At 7;7 years the Spanish version of the Inventory for Client and Agency Planning (ICAP) ^10^ was administered in order to evaluate her behavioral disturbances. The obtained scores were indicative of moderately-serious asocial maladaptive behavior (uncooperative behavior) and of moderately-serious externalized maladaptive behavior (disruptive behavior and hurtful to others). The ICAP scores were in line with the scores obtained in the Battelle subtest for personal-social abilities, which were suggestive of problems for correctly interacting with her peers and for establishing emotional links, and ultimately, of uncooperative and disruptive behavior. Because of the difficulties experienced by the proband she has attended a special education unit since the age of 6. From 6 years until present she has been learning Spanish Sign Language, which was thought to aid in improving her communication abilities.

**Figure 1.**
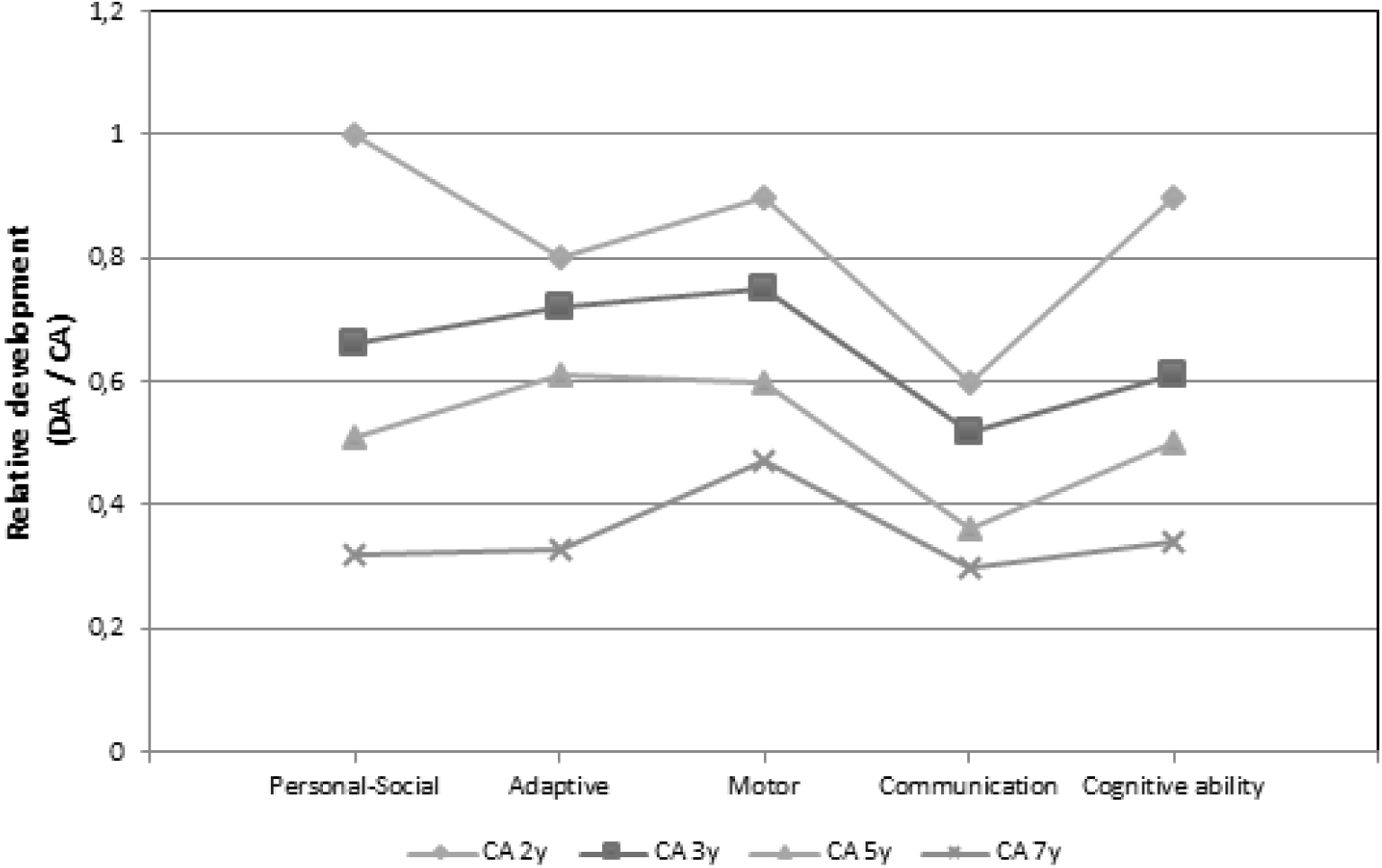
Proband’s developmental profile according to the Battelle Developmental Inventories. In order to make more reliable comparisons, the resulting scores are shown as relative values referred to the expected scores according to the chronological age of the child. Abbreviations: DA, developmental age; CA, chronological age.

At 7;7 years, her language problems were assessed in detail. Language comprehension abilities were assessed with the Spanish version of the Peabody Picture Vocabulary Test, Third Edition (PPVT-3) ^11^, whereas language production was assessed with the Prueba de Lenguaje Oral de Navarra-Revisada [Navarra Oral Language Test, Revised] (PLON-R) ^12^, and by means of a natural speech sample. In the PPVT-3 the proband scored 47 points (3;9 years below expected). In the PLON-R the proband scored 4 years below her TD peers. In her discourse she exhibited phonological deformations that are normally found in younger children, including reductions of consonant clusters, deletions of unstressed syllables, assimilations, and substitutions. Nonetheless, the child was able to properly articulate all the Spanish phonemes separately. On the morphological and syntactic side, she had problems for correctly repeating the target sentences provided by the experimenter. Hence, in the elicited production tasks of the PLON-R, the proband usually made use of simpler syntactic structures (mostly, list of words). Similarly, she correctly produced terms for colors, spatial relations, body parts, and simple actions, but only when the test aimed to test younger children (3 years) was administered. Overall, the semantic content of the proband’s sentences was relatively spared, but the sentence structure was largely impaired. The analysis of spontaneous speech samples generated by the child supported the view that her language was structurally simpler than that of her TD peers. Her discourse mostly consisted of simple sentences and questions about her surrounding environment or favorite topics. Verbs and functional words were usually omitted. Sentence modality was correctly marked by prosodic cues (i.e. voice inflections). Linguistically, the child’s impulsive behavior and lack of self-control resulted in logorrhea (mostly, in the form of repetitive fragments of speech), but no signs of echolalia were observed. Interestingly, the child used to ‘sing without words’ privately (i.e. generate an unstructured melodic emission) as a form of self-stimulation whenever she was compelled to fulfill tasks that were difficult to achieve (e.g. those encompassing the evaluation tests). Language abilities with sign language were further evaluated. They did not differ from the abilities exhibited in the oral domain, in spite of her slight hearing loss. Comprehension clearly surpassed production. Signs were poorly articulated.

### Molecular Cytogenetic Analysis

Routine molecular cytogenetic analyses were performed at age 3;5. Polymerase chain reaction (PCR) analysis of the *FMR1* fragile site was normal. Fluorescent in situ hybridization (FISH) analyses of the loci of *SNRPN* and *UBE3A* were also normal. No major chromosomal rearrangements were observed in the karyotype analysis. Because the analysis did not discard the presence of low frequency mosaicisms and/or cryptic chromosomal alterations, a comparative genomic hybridization array (array-CGH) (Cytochip Oligo ISCA 60K) was subsequently performed (DLRS value = 0.15; data were analyzed by ADM-2 (Agilent Technologies): threshold = 6.0; aberrant regions had more than 5 consecutive probes). The array identified five copy number variants (CNVs): one large deletion within region 7q32.1-q33 (chr7:127109685-132492196, hg18), two small deletions within regions 8p23.1 (chr8:7156900-7359099 hg18) and 15q13.1 (chr15:26215673-26884937), respectively, and two microinsertions within region Xp22.33 (chrX:17245-102434 and chrX:964441-965024, respectively) (figure 2). The parents’ karyotypes were normal and the arrays did not identified any CNVs in any of them.

**Figure 2.**
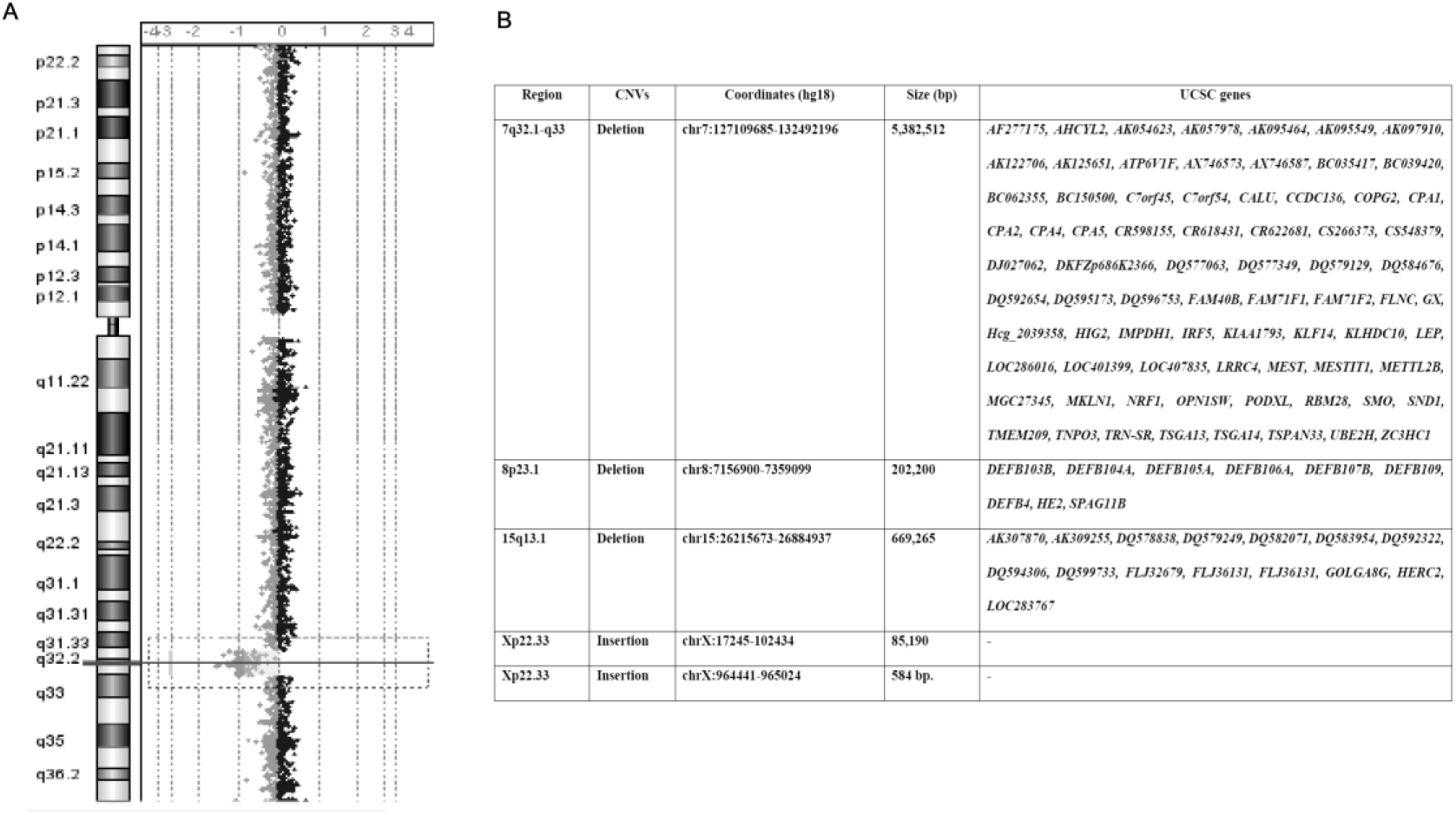
Array-CGH and deleted genes in the proband. A. Array-CGH of the proband’s chromosome 7 showing the microdeletion in 7q32.1-q33. B. Summary table of the genes deleted in the proband.

## DISCUSSION

We have described a child with severe language impairment (particularly in the oral domain) and cognitive and behavioural disturbances, and with several de novo chromosomal rearrangements that have mostly impacted chromosome 7. Several candidates for language and cognitive disorders are located within the regions deleted in the proband. With regards to the language deficits exhibited by our proband, we wish highlight that the fragment deleted in 7q32.1-q33 contains several targets of FOXP2, including *CALU*, *IMPDH1*, *LEP*, *MEST*, *OPN1SW*, and *RBM28* ^13,14^. *FOXP2* is a well-known language-related gene that contributes to the development of cortico-thalamic-striatal circuits involved in procedural learning and whose mutation gives rise to speech and language problems ^3,4^. Interestingly, *CALU* codes for a calumenin, a calcium-binding protein of the endoplasmic reticulum. The gene is expressed highly during early development in different brain regions, particularly in migrating neurons ^15^. Mutations in *RBM28* affect ribosome biogenesis and the structure of the rough endoplasmic reticulum, and give rise to microcephaly, moderate to severe mental retardation, and progressive motor deterioration resulting from upper and lower motor dysfunction ^16^. Within the deleted fragment in chromosome 7 we have found as well two targets (*CPA4* and *MKLN1*) of another gene that we believe important for the development and the evolution of language, namely, *RUNX2. RUNX2* encodes an osteogenic factor that controls the closure of cranial sutures and several aspects of brain growth, and has been related to most of the changes that brought about our more globular brain(case) and our species-specific mode of cognition, including language ^17,18^.

Additionally, several of the genes deleted in the proband belong to the autism susceptibility locus on 7q32 (AUTS9). These include both imprinted (*MEST*, *CPA4*, *CPA5, KLF14*) and non-imprinted (*COPG2, TSGA14*) genes. Interestingly, *COPG2* expression is downregulated in the human cortex compared to the chimpanzee, supporting a role for this gene in the development of a ‘social brain’ (according to ^2^). Similarly, point mutations in *TSGA14* ^19^ and *UBE2H* ^20^ have been identified in families with autism spectrum disorders. The latter encodes an E2 (ubiquitin-conjugating) enzyme involved in neural development (intriguingly, another member of this system, *UBE3A*, causes Angelman syndrome). Changes in the expression levels of imprinted genes within this region have been related as well to Russell-Silver Syndrome ^21^, a condition involving intrauterine growth retardation, poor postnatal growth, and craniofacial malformations ^22^.

Two other genes of interest also deleted in the proband are *PODXL* and *NRF1*. Both of them play a role in brain development and function. The former encodes a polysialylated neural adhesion protein involved in neurite growth, neuron branching, and axonal fasciculation; in mice the loss of *Podxl* function results in a reduction of the number of synapses in the central nervous system and in the neuromuscular system ^23^. Regarding *NRF1*, it encodes a transcription factor that regulates many nuclear genes that are essential for mitochondrial function; this gene seems to be relevant to the pathogenesis of neurodegenerative diseases via perturbation of mitochondrial and extra-mitochondrial functions ^24^.

Outside the region deleted in chromosome 7, we have found one gene only that may contribute to the cognitive and linguistic profile of our proband: *HERC2*. According to ^25^, mutations in *HERC2* affect E3 ubiquitin ligase activity, also resembling the pathophysiologic mechanism underlying Angelman syndrome. This gene is a candidate for autosomal recessive mental retardation-38 (MRT38), a condition involving motor, speech, adaptive, and social delay, autistic features, aggression, impulsivity, and distractibility ^25^.

Finally, we wish briefly mention some other genes that are located within the region deleted in chromosome 7 and that may contribute to some of the proband’s dysfunctions in other domains. Specifically, the loss of *FLNC*, a gene encoding filamin C and related to myofibrillar myopathy ^26^, may cause the motor problems exhibited by the child. The hemyzigosity of *SMO* and *UBE2H* (two genes expressed in the choclea), and of *MIR96* (related to deafness ^27,28^) may account for her hearing problems. Interestingly as well, most of the genes contained in the deleted region in chromosome 8 encode defensines, some of which (specifically DEFB4) have been linked to an enhanced susceptibility to infectious disease affecting the ear (particularly, otitis media) ^29^.

In conclusion, although the exact genetic cause of the language and cognitive impairment exhibited by our proband remains to be fully elucidated, we believe that the hemyzigosity of the genes discussed above may explain most of her deficits and clinical problems. We expect that this case (and the genes we highlight in this paper) contribute as well to a better understanding of the genetic underpinnings of the human faculty for language, particularly, of the genes located in 7q31-q33.

## ETHICS, CONSENT, AND PERMISSIONS

Ethics approval for this research was granted by the Comité Ético del Hospital Universitario “Reina Sofía”. Written informed consent was obtained from the proband’s parents for publication of this case report and of any accompanying tables and images. A copy of the written consent is available for review by the Editor-in-Chief of this journal. The authors declare that none of them have any competing interests.

## ACKNOWLEDGMENTS

We would like to thank the proband and her family for their participation in this research.

### FUNDING ACKNOWLEDGMENTS

This work was supported by the Spanish Ministry of Economy and Competitiveness (grant numbers FFI-2013-43823-P and FFI2014-61888-EXP to Antonio Benítez-Burraco).

## REFERENCES

1. Benítez-Burraco A. Genetics of language: Roots of specific language deficits. In: Boeckx C, Grohmann KK, editors. The Cambridge Handbook of Biolinguistics. Cambridge: Cambridge University Press; 2013:375–412.

2. Schneider E, Jensen LR, Farcas R, Kondova I, Bontrop RE, Navarro B, et al. A high density of human communication-associated genes in chromosome 7q31-q36: differential expression in human and non-human primate cortices. Cytogenet Genome Res. 2012; 136:97–106

3. Fisher SE, Scharff C. FOXP2 as a molecular window into speech and language. Trends Genet. 2009; 25:166–77.

4. Schreiweis C, Bornschein U, Burguière E, Kerimoglu C, Schreiter S, Dannemann M et al. Humanized Foxp2 accelerates learning by enhancing transitions from declarative to procedural performance. Proc. Natl. Acad. Sci. U.S.A. 2014; 111:14253–14258.

5. Kaminen N, Hannula-Jouppi K, Kestilä M, Lahermo P, Muller K, Kaaranen M, et al. A genome scan for developmental dyslexia confirms linkage to chromosome 2p11 and suggests a new locus on 7q32. J Med Genet. 2003; 40:340–345.

6. Peter B, Matsushita M, Raskind WH. Motor sequencing deficit as an endophenotype of speech sound disorder: a genome-wide linkage analysis in a multigenerational family. Psychiatr Genet. 2012; 22:226–234.

7. Beyer KS, Klauck SM, Wiemann S, Poustka A. Construction of a physical map of an autism susceptibility region in 7q32.3-q33. Gene 2001; 272:85–91.

8. Bonora E, Bacchelli E, Levy ER, Blasi F, Marlow A, Monaco AP, et al. Mutation screening and imprinting analysis of four candidate genes for autism in the 7q32 region. Mol Psychiatry 2002; 7:289–301.

9. De La Cruz López MV, González Criado M. Adaptación española del inventario de Desarrollo Battelle. Madrid: TEA, 2011

10. Montero D. Evaluación de la conducta adaptativa en personas con discapacidades. Adaptación y validación del ICAP. Bilbao: Mensajero; 1996.

11. Dunn L, Dunn LM, Arribas D. Peabody, test de vocabulario en imágenes. Madrid: Tea Ediciones; 2006.

12. Aguinagua G, Armentia M, Fraile A, Olangua P, Uriz N. PLON.-R Prueba de Lenguaje Oral de Navarra. Madrid: TEA; 2004.

13. Vernes SC, Oliver PL, Spiteri E, Lockstone HE, Puliyadi R, Taylor JM, et al. Foxp2 regulates gene networks implicated in neurite outgrowth in the developing brain. PLoS Genet. 2011; 7:e1002145.

14. Vernes SC, Spiteri E, Nicod J, Groszer M, Taylor JM, Davies KE et al. High-throughput analysis of promoter occupancy reveals direct neural targets of FOXP2, a gene mutated in speech and language disorders. Am. J. Hum. Genet. 2007; 81:1232–1250.

15. Vasiljevic M, Heisler FF, Hausrat TJ, Fehr S, Milenkovic I, Kneussel M, et al. Spatio-temporal expression analysis of the calcium-binding protein calumenin in the rodent brain. Neuroscience 2012; 202:29–41.

16. Nousbeck J, Spiegel R, Ishida-Yamamoto A, Indelman M, Shani-Adir A, Adir N, et al. Alopecia, neurological defects, and endocrinopathy syndrome caused by decreased expression of RBM28, a nucleolar protein associated with ribosome biogenesis. Am J Hum Genet. 2008; 82:1114–1121.

17. Boeckx C, Benítez-Burraco A. The shape of the human language-ready brain. Front. Psychol. 2014a; 5:282.

18. Benítez-Burraco A, Boeckx C. Possible functional links among brain-and skull-related genes selected in modern humans. Front. Psychol. 2015; 6:794

19. Korvatska O, Estes A, Munson J, Dawson G, Bekris LM, Kohen R, et al. Mutations in the TSGA14 gene in families with autism spectrum disorders. Am J Med Genet B Neuropsychiatr Genet. 2011; 156B:303–311.

20. Vourc’h P, Martin I, Bonnet-Brilhault F, Marouillat S, Barthélémy C, Pierre Müh J, et al. Mutation screening and association study of the UBE2H gene on chromosome 7q32 in autistic disorder. Psychiatr Genet. 2003; 13:221–225.

21. Penaherrera MS, Weindler S, Van Allen MI, Yong S-L, Metzger DL, McGillivray B et al. Methylation profiling in individuals with Russell-Silver syndrome. Am J Med Genet 2010; 152:347–355

22. Price SM, Stanhope R, Garrett C, Preece MA, Trembath RC. The spectrum of Silver-Russell syndrome: a clinical and molecular genetic study and new diagnostic criteria. J Med Genet 1999; 36:837–842

23. Vitureira N, Andrés R, Pérez-Martínez E, Martínez A, Bribián A, Blasi J, et al. Podocalyxin is a novel polysialylated neural adhesion protein with multiple roles in neural development and synapse formation. PLoS One. 2010; 5:e12003.

24. Satoh J, Kawana N, Yamamoto Y. Pathway Analysis of ChIP-Seq-Based NRF1 Target Genes Suggests a Logical Hypothesis of their Involvement in the Pathogenesis of Neurodegenerative Diseases. Gene Regul Syst Bio. 2013; 7:139–152.

25. Puffenberger EG, Jinks RN, Wang H, Xin B, Fiorentini C, Sherman EA et al. A homozygous missense mutation in HERC2 associated with global developmental delay and autism spectrum disorder. Hum. Mutat. 2012; 33:1639–1646

26. Tasca G, Odgerel Z, Monforte M, Aurino S, Clarke NF, Waddell LB, et al. Novel FLNC mutation in a patient with myofibrillar myopathy in combination with late-onset cerebellar ataxia. Muscle Nerve 2012; 46:275–282.

27. Friedman LM, Avraham KB. MicroRNAs and epigenetic regulation in the mammalian inner ear: implications for deafness. Mamm Genome 2009; 20:581–603.

28. Soldà G, Robusto M, Primignani P, Castorina P, Benzoni E, Cesarani A et al. A novel mutation within the MIR96 gene causes non-syndromic inherited hearing loss in an Italian family by altering pre-miRNA processing. Hum Mol Genet 2012; 21:577–585.

29. Jones EA, Kananurak A, Bevins CL, Hollox EJ, Bakaletz LO. Copy number variation of the beta defensin gene cluster on chromosome 8p influences the bacterial microbiota within the nasopharynx of otitis-prone children. PLoS One 2014; 9:e98269.

